# Healthy dietary choices involve prefrontal mechanisms associated with long-term reward maximization but not cognitive control

**DOI:** 10.1101/2024.02.15.580403

**Authors:** Ai Takehana, Daiki Tanaka, Mariko Arai, Yoshiki Hattori, Takaaki Yoshimoto, Teppei Matsui, Norihiro Sadato, Junichi Chikazoe, Koji Jimura

**Affiliations:** Department of Informatics, Gunma University, Maebashi, Japan; Faculty of Biological and Environmental Sciences, University of Helsinki, Helsinki, Finland; Department of Biosciences and Informatics, Keio University, Yokohama, Japan; Supportive Center for Brain Research, National Institute for Physiological Sciences, Okazaki, Japan; Araya Inc., Tokyo, Japan; Graduate School of Brain Science, Doshisha University, Kyotanabe, Japan; Research Organization of Science and Technology, Ritsumeikan University, Kusatsu, Japan

**Author notes:** Correspondence should be addressed to: Koji Jimura, Ph.D. Department of Informatics, Gunma University, 4-2 Aramaki-machi Maebashi, 371-8510, Japan.

## Abstract

Taste and health are critical factors to be considered when choosing foods. Prioritizing healthiness over tastiness requires self-control. It has also been suggested that self-control is guided by cognitive control. We then hypothesized that neural mechanisms underlying healthy food choice are associated with both self-control and cognitive control. Human participants performed a food choice task and a working memory (WM) task during functional MRI scanning. Their degree of self-control was assessed behaviorally by the value discount of delayed monetary rewards in intertemporal choice. Prioritizing healthiness in food choice was associated with greater activity in the superior, dorsolateral, and medial prefrontal cortices. Importantly, the prefrontal activity was greater in individuals with smaller delay discounting (i.e., high self-control) who preferred a delayed larger reward to an immediate smaller reward in intertemporal choice. On the other hand, WM activity did not show a correlation with delay discounting or food choice activity, which was further supported by supplementary results that analyzed data from the Human Connectome Project. Our results suggest that the prefrontal cortex plays a critical role in healthy food choice, which requires self-control, but not cognitive control, for maximization of reward attainments in a remote future.

## Introduction

Taste and healthiness are significant factors in choosing foods (Hare et al., 2011). Making appropriate food choices is essential to ensure good health. However, humans often impulsively choose unhealthy foods by prioritizing those that taste good (e.g., high-calorie foods) (van der Laan et al. 2014). Such impulsive food choice is attributable to the distinct reward types inherent in diets: pleasant taste provides immediate satisfaction, but achieving good health takes a longer time (Fishbach et al. 2003). Thus, food choice often imposes a tradeoff between short- and long-term rewards (i.e., taste and healthiness, respectively).

Previous studies have suggested that healthy food choice involves self-control that prioritizes a long-term reward over a short-term reward (Hare et al. 2011; van der Laan *et al*. 2014; Blechert et al. 2016). Thus, self-controlled individuals are more likely to choose healthy foods, while impulsive individuals may tend to prioritize favorable taste.

A tradeoff between short- and long-term rewards has been well examined in intertemporal decision-making. In a standard choice situation, agents make a choice between a smaller amount of a reward available soon and a larger amount of a reward delivered after a long delay (Keeney and Raiffa 1993; Green and Myerson 2004). It is well known that agents discount the values of rewards available in a remote future. This phenomenon, known as delay discounting, is measured as a representative index reflecting the degree of self-control (Madden and Bickel 2009; Peters and Büchel 2011). Smaller delay discounting indicates a preference for larger delayed rewards and reflects strong self-control (Kirby et al. 1999; Luo et al. 2009; Figner et al. 2010). By contrast, a preference for smaller immediate rewards is characterized as impulsive and presents greater delay discounting (Madden and Bickel 2009).

Notably, the food choice and standard intertemporal choice both involve a tradeoff between an immediate reward (i.e., food choice: taste, intertemporal choice: smaller reward) and a temporally remote reward (i.e., food choice: health, intertemporal choice: larger reward). Although the source of tradeoff is not fully matched (food choice: taste and health; intertemporal choice: amount), choice preference has been explained along an axis from self-control to impulsivity in both situations (Keeney and Raiffa 1993; Kirby *et al*. 1999; Green and Myerson 2004; Hare et al. 2009; Luo *et al*. 2009; Madden and Bickel 2009; Figner *et al*. 2010; Peters and Büchel 2011; van der Laan *et al*. 2014; Blechert *et al*. 2016). Impulsivity in food choice often induces excessive intake of tasty, high-calorie foods, which leads to obesity (Davis et al. 2004; Davis et al. 2010; Berthoud and Zheng 2012; Schiff et al. 2016; Spetter et al. 2017).

Interestingly, individuals with obesity showed greater delay discounting in intertemporal choice for monetary rewards (Davis *et al*. 2004; Davis *et al*. 2010; Stoeckel et al. 2013; Lawyer et al. 2015; Zhang et al. 2022). These observations suggest that strong impulsivity in food choice is associated with strong impulsivity as indexed by the degree of delay discounting.

Neuroimaging studies of food choice have shown that self-controlled decision-making is associated with strong activations of the dorsolateral prefrontal cortex (dlPFC) and the ventromedial prefrontal cortex (vmPFC) (Hare *et al*. 2009; Maier et al. 2015; Spetter *et al*. 2017). Interestingly, it has also been shown that self-control in intertemporal choice involves the dlPFC (McClure et al. 2004; Figner *et al*. 2010), and reward values are represented in the vmPFC (Kable and Glimcher 2007). These neuroimaging studies collectively suggest that the involvement of the dlPFC and vmPFC in self-controlled food choice reflects self-control in intertemporal choice. As such, these affinities regarding theoretical accounts and empirical evidence between food choice and intertemporal choice suggest a link between them.

Cognitive control helps to achieve a behavioral goal and involves orientation of attention, updating goal-relevant information, and withdrawal of inappropriate behavior (Miller and Cohen 2001). It is thought that cognitive control guides self-control in decision-making (Duckworth and Seligman 2005; Frederick 2005). A representative, central function of cognitive control is working memory (WM) that enables encoding, active maintenance, and updating of goal-relevant information implicated in the dlPFC (D’Esposito and Postle 2015). Greater WM activity in the dlPFC is associated with higher general intelligence (Gray et al. 2003), and higher intelligence is also associated with smaller delay discounting in intertemporal choice (Shamosh and Gray 2008). It is thus reasonable that strong WM-related activity is associated with smaller delay discounting (Hinson et al. 2003; Shamosh et al. 2008).

Value-based decision-making often requires integrative computation of choice information. In decision-making where choice information involves a high-dimensional complexity of multiple factors, fronto-parietal regions showed greater activity, and the activity reflects the engagement of WM (Matsui et al. 2022). Additionally, in a situation where the disparity in reward values of the presented options is smaller, the decision-making becomes more difficult and thus needs elaborate computation of choice information. Such difficult decision-making also involves prefrontal mechanisms for WM (Jimura et al. 2018).

Given the close relationships between WM and delay discounting and between delay discounting and healthy food choice, we hypothesized that self-control in healthy food choice is linked to both high self-control (smaller delay discounting in intertemporal choice) and high cognitive control.

To test this hypothesis, we performed a series of experiments. Human participants first subjectively rated the taste and healthiness of food items (Fig. 1A). They then chose between the food items during functional MRI scanning (Fig. 1B), which allowed us to identify brain regions involved in prioritizing healthiness over taste. The participants’ degree of self-control was also assessed based on an intertemporal choice task using a delayed monetary reward (Fig. 1C). Finally, they were scanned while performing a WM task (Fig. 1D). Our hypothesis predicted that:

1. the dlPFC and vmPFC show prominent activity during self-controlled food choice, and their activity is enhanced in individuals showing high self-control (smaller delay discounting) in intertemporal choice,
2. brain regions involved in healthy food choice are also involved in WM, and WM-related activity is enhanced in individuals with strong activity in self-controlled food choice, and
3. WM-related activity is enhanced in individuals with high self-control in intertemporal choice.

**Figure 1.**
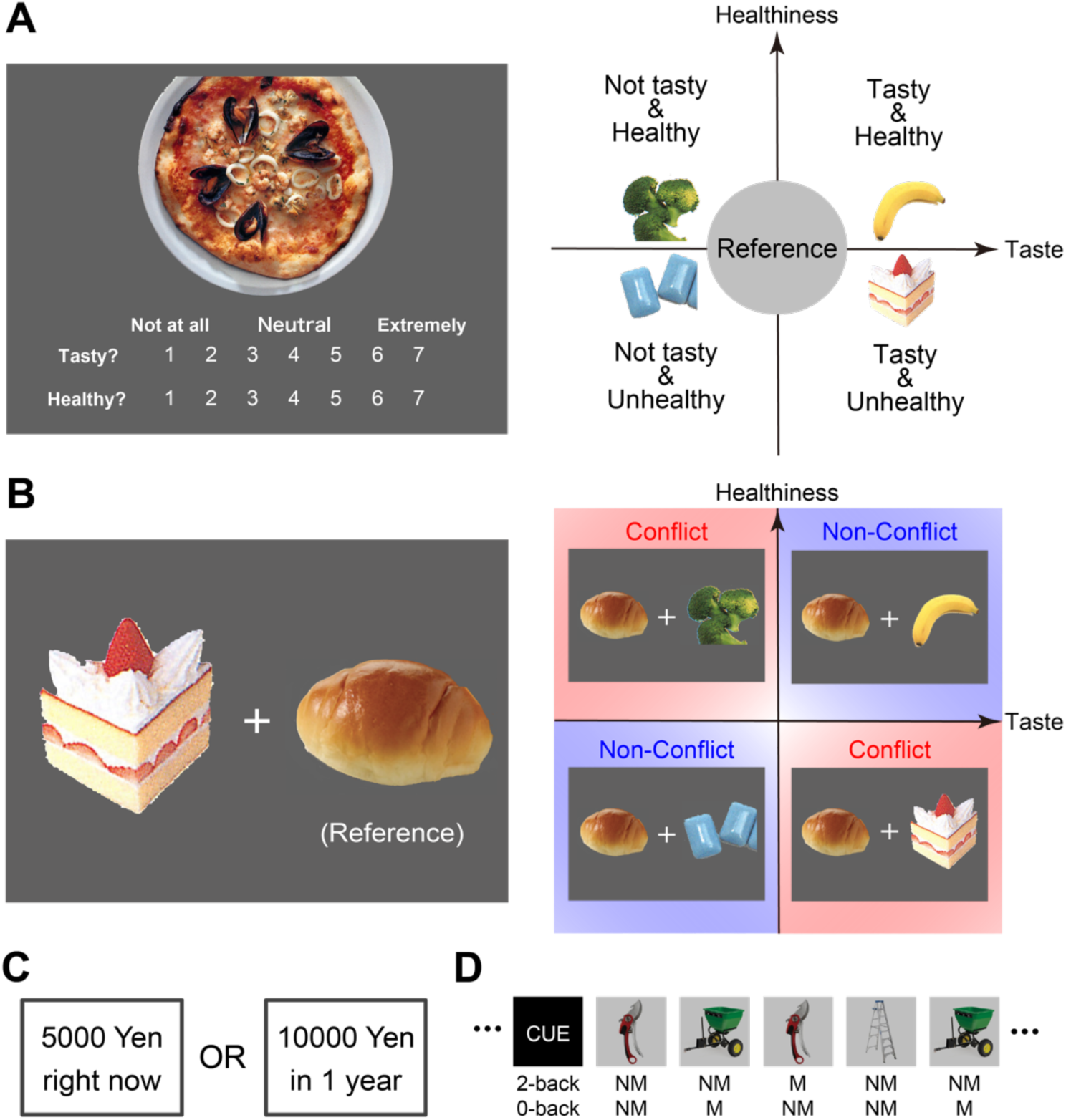
Study design. ***A***, Prior to fMRI scanning, participants rated the taste and healthiness of food items on a seven-point scale (*left*). *Right*) Based on the rating, each food item was classified as Tasty– Healthy (*upper right* quadrant; e.g., banana), Not tasty–Healthy (*upper left* quadrant; e.g., broccoli), Not tasty–Unhealthy (*bottom left* quadrant; e.g., chewing gum), Tasty–Unhealthy (*bottom right* quadrant; e.g., strawberry shortcake), or as a Reference item (*center*). ***B***, During fMRI scanning, participants made a choice between two food items (*left*). One item belonged to one of the Tasty–Healthy, Not tasty–Healthy, Not tasty–Unhealthy, Tasty–Unhealthy categories, and the other item was a reference item (*right*; e.g., bread). In the Not tasty–Heathy and Tasty–Unhealthy trials, there was a tradeoff between taste and healthiness (conflict trials), whereas this tradeoff was absent in the Tasty–Healthy and Not tasty– Unhealthy trials (non-conflict trials). ***C***, Outside the fMRI scanner, participants performed an intertemporal choice task with a hypothetical monetary reward to measure their degree of self-control. Procedures were based on those used in the Human Connectome Project (HCP). ***D***, During fMRI scanning, participants performed an N-back working memory task that was used in the HCP. M: match; MN: non-match.

Collectively, these behavioral and neuroimaging experiments revealed relationships between 1) delay discounting and WM activity and 2) food choice activity and WM activity.

## Materials and Methods

### Participants

Written informed consent was obtained from 49 young, healthy participants (age range: 18-28; 23 females). Experimental and analytical procedures were approved by the institutional review boards of Gunma University, Keio University, and the National Institute for Physiological Sciences. The sample size was determined prior to the experiment based on power analysis (α = 0.05, β = 0.95, η^2^ = 0.5).

### Food rating and choice tasks: rating healthiness and taste

#### Stimuli

A list of 117 food items was compiled based on books and websites. The food items included vegetables, fruits, main courses, fast foods, snacks, and sweets. All visual stimuli were presented using E-Prime (Psychology Software Tools, Sharpsburg PA, USA).

### Rating task procedures

In each trial, the image of one food item was presented at the center of a computer screen (Fig. 1A *left*), and participants subjectively rated the healthiness and taste of the item on a 7-point scale. The taste rating scale ranged from “1” (not tasty) to “7” (very tasty). The health rating scale ranged from “1” (very unhealthy) to “7” (very healthy). Participants responded by pressing a number key (from “0” to “7”) on the computer keyboard. If they were unfamiliar with the food item, had never eaten it, or had an allergy to it, they were instructed to press “0.” When rating taste, participants were instructed to consider taste only, and not healthiness; conversely, when rating healthiness they were instructed not to consider taste. They were also instructed to balance their rates using the entire range of the scale, not using a particular range.

After the ratings were complete, the items were classified into five types according to each participant’s ratings (Fig. 1A *right*). The reference food type consisted of items with median ratings for both taste and healthiness. The other four types were defined as follows: 1) Tasty–Healthy (taste > median, and healthiness > median), 2) Not tasty–Healthy (taste < median, and healthiness > median), 3) Not tasty–Unhealthy (taste < median, and healthiness < median), and 4) Tasty–Unhealthy (taste > median, and healthiness < median). The food items that the participants rated 0 (i.e., they were unfamiliar with, had not eaten, or had an allergy to them) were not included in the reference type or the other four types.

### Choice task procedures

During fMRI scanning, participants performed a food choice task. In each trial, two food items were presented on the left and right sides of the screen: one of these was a reference item, and the other (the target item) belonged to one of the other four food types (Tasty–Healthy, Not tasty–Healthy, Not tasty–Unhealthy, or Tasty–Unhealthy; Fig. 1B). The two items were randomly placed. Participants were instructed to choose the item that they would prefer to eat, and then to press the corresponding button with their right thumb. Choice items disappeared when participants made a button response, followed by a 2-s inter-trial interval.

Each functional run consisted of 40 trials (10 trials for each of the four target item types) and lasted 270 s. Participants completed three functional runs, consisting of 120 choice trials in total.

### Intertemporal decision-making task

To measure the degree of self-control of each participant, participants performed an intertemporal choice task for hypothetical money prior to fMRI scanning (Fig. 1C). Task procedures were described in the previous studies (Jimura et al. 2009; Jimura et al. 2011; Jimura et al. 2013; Jimura *et al*. 2018). Importantly, this procedure was similar to that used in the HCP (Estle et al. 2006; Green et al. 2007; Barch et al. 2013).

Specifically, in each trial, two choices were presented on the screen: a larger reward at a later point in time, and a smaller reward available immediately (Fig. 1C). Participants were instructed to think of the choices as if they were real and to choose the one that they preferred. Choices were presented to the left and right of a central fixation point, and the positions of the immediate and delayed rewards varied randomly with each trial. Participants indicated their preference by pressing either the “1” (left) or “2” (right) key on the computer keyboard.

Participants made choices regarding delays of six different durations (1 week, 1 month, 3 months, 6 months, 1 year, 3 years); the reward after each delay was the same in each case (10,000 yen). In the first trial, the immediate reward was 5,000 yen. The reward amounts in subsequent trials were adjusted depending on each participant’s selection. Specifically, if the participant chose the immediate reward, then the immediate reward amount was decreased by half for the second choice (2,500 yen); if the participant chose the delayed reward, then the immediate reward amount was increased by half (7,500 yen). The change in the immediate reward amount in each subsequent choice was 1/2 that in the preceding choice. More specifically, in the second choice, the immediate reward was increased or decreased by 2,500 yen, in the third choice by 1,250 yen, in the fourth choice by 625 yen. After the four choices, then the subjective value of the delayed reward was estimated to be equal to 312.5 yen more or less than the immediate reward amount available in the fourth trial, depending on whether the delayed or immediate reward had been chosen in that trial.

### WM task

Participants also performed an N-back WM task during fMRI scanning (Fig. 1D). The task was identical to the one used in the HCP (http://www.humanconnectome.org; Barch et al. 2013). We used E-Prime scripts provided from the Connectome DB (https://db.humanconnectome.org/).

Prior to each task block, a cue stimulus was presented at the center of the screen, indicating the task condition (2-back or 0-back). For the 0-back condition, a target item was presented simultaneously. The cue stimulus presentation was followed by the presentation of 10 pictures, with each presented for 2.5 s, followed by a 0.5-s intertrial interval. The 10 pictures in a single block were drawn from one of four categories (body parts, faces, places, or tools). One functional run involved eight task blocks (two WM conditions x four picture categories). Four 15-s fixation task blocks were pseudorandomly inserted among the task blocks. Each participant performed two functional runs.

### Imaging procedures

MRI scanning was performed using a 3-T MRI scanner (Siemens Verio, Germany) with a 32-channel head coil. Functional images were acquired using a multiband acceleration echo-planar imaging sequence [repetition time (TR): 800 msec; echo time (TE): 30 msec; flip angle (FA): 45 deg; 72 slices; slice thickness: 2 mm; in-plane resolution: 2 x 2 mm; multiband factor: 8]. For the food choice task, one functional run lasted 270 s with 338 volume acquisitions, while for the working memory task, one functional run lasted 300 s with 365 volume acquisitions. The first 10 volumes in each functional run were discarded to take into account the time required for equilibrium of longitudinal magnetization. High-resolution anatomical images were acquired using an MP-RAGE T1-weighted sequence [TR: 1900 msec; TE = 2.52 msec; FA: 9 deg; 176 slices; slice thickness: 1 mm; in-plane resolution: 1 x 1 mm^2^].

### Imaging preprocessing

MRI data were analyzed using SPM12 software (http://www.fil.ion.ucl.ac.uk/spm/ software/spm12/). All functional images were first temporally aligned across volumes and runs, and then each anatomical image was coregistered to the mean of the corresponding functional images. The functional images were spatially normalized to a standard MNI template with normalization parameters estimated from the corresponding anatomical image. The images were then resampled into 2-mm isotropic voxels, and spatially smoothed with a 6-mm full-width at half-maximum (FWHM) Gaussian kernel.

### Imaging analysis: first-level analysis

A general linear model (GLM) (Worsley and Friston 1995) approach was used to estimate parameter values for task events.

For the food choice task, the trials of interest were those in which 1) the rating for taste or healthiness was greater than that of the reference item and 2) the other rating (healthiness or taste) was smaller than that of the reference item (conflict trials; Not tasty–Healthy and Tasty–Unhealthy trials; Fig 1B *right*). In these trials, participants’ choices were parametrically coded. Specifically, the conflict trials in which a high-health, low-taste item was chosen (i.e., the target item in Not tasty–Healthy trials and the reference item in Tasty-Unhealthy trials) were coded as 1, while those in which the low-health, high-taste item was chosen were coded as −1. The parametric regressor was then z-scored (mean-centered and divided by its standard deviation). Tasty–Healthy and Not tasty–Unhealthy trials were separately coded as trials that were not of interest.

These task events were time-locked to the onset of food item presentation and then convolved with a canonical hemodynamic response function (HRF) implemented in SPM. Additionally, six-axis head movement parameters, white-matter signals, CSF signals, and whole-brain signals were included in the GLM as nuisance effects. Then, parameters were estimated for each voxel across the whole brain.

A separate GLM analysis was performed for the WM task. Each of the eight block types (two WM conditions for four picture categories) was separately coded and then convolved with a canonical HRF implemented in SPM. Similarly to the food choice task, six-axis head movement parameters, white-matter signals, CSF signals, and whole brain signals were included in the GLM as nuisance effects. Then, parameters were estimated for each voxel across the whole brain.

### Group-level analysis

For the food task, maps of the parameter estimate for the parametric effect of the choice in the food choice task were collected from all participants. For the WM task, maps of the parameter estimate were first contrasted between 2-back and 0-back conditions for individual participants, and the contrast maps were collected from all participants.

These maps were then subjected to group-level one-sample t-tests based on permutation methods (5000 permutations) implemented in *randomise* in the FSL suite (Eklund et al. 2016) (http://fmrib.ox.ac.uk/fsl/). Voxel clusters were identified using a voxel-wise uncorrected threshold of P < .01 for the food choice task and P < .001 for the WM task, and the identified voxel clusters were tested for significance with a threshold of P < .05 corrected by the family-wise error (FWE) rate across the whole brain.

This procedure was empirically validated to appropriately control the false-positive rate when correcting P-values across the whole brain (Eklund et al., 2016). This procedure was also robust against the levels of the cluster-defining threshold (Eklund et al., 2016). The peaks of significant clusters were then identified and listed in tables. If multiple peaks were identified within 12 mm, the most significant peak was kept.

### Assessment of self-control

Individual differences in delay discounting were quantified by calculating the area under the curve (AuC) of individuals’ discounting plots (Myerson et al. 2001; Sellitto et al. 2010; Jimura *et al*. 2011; Jimura *et al*. 2013; Jimura *et al*. 2018; Tanaka et al. 2020). The AuC represents the area under the observed subjective values, and was calculated as the sum of the trapezoidal areas under the indifference points normalized by the amount and delay (Myerson et al. 2001). It has been argued that the AuC is a valid measure of delay discounting for use in individual difference analyses, because it is theoretically neutral and also psychometrically reliable (Myerson et al. 2001). Participants with a higher AuC are characterized as self-controlled, and those with a low AuC show and are characterized as impulsive.

Importantly, the AuC measures for the delay discounting curve were also used to represent the degree of self-control in the HCP (Barch et al. 2013), allowing for compatible of self-control assessment between the current study and the HCP.

### Exploratory correlation analysis of AuC and brain activity maps

To examine whether brain activity during the food and WM tasks was modulated by individuals’ degree of self-control, correlational imaging analysis was performed (Jimura *et al*. 2013; Tanaka *et al*. 2020; Shintaki et al. 2022). Voxel-wise correlation coefficients were calculated for the activity maps of the parametric choice effect in the food choice task and 2-back vs. 0-back contrast in the WM task, and corresponding t-maps were created. Voxel clusters were then identified using a voxel-wise uncorrected threshold of P < .05, and the identified voxel clusters were tested for significance with a threshold of P < .05 corrected by the FWE rate across the whole brain.

### Complementary analysis based on HCP data

One notable design feature of the current study is that we adopted the self-control assessment and WM task procedures used in the HCP, allowing us to perform complementary analysis utilizing the large HCP sample. From the Connectome DB, we collected event onsets and preprocessed 4D timeseries of fMRI images obtained during the WM task, as well as AuC measures for the delay-discounting task (N = 1075). For compatibility between the current study and the HCP, we performed first-level GLM estimations for the WM task using SPM. Then, we performed group-level analysis using procedures identical to those utilized for the data collected in the current experiment (see the “Group-level analysis” section above).

## Results

### Behavioral results

#### Food choice task

In the food choice task (Fig. 1B), we expected that participants would choose food that was both healthy and tasty over the reference item, when such a choice was available. Conversely, participants would choose the reference item when the other food was both unhealthy and not tasty. As expected, the rate of choosing the reference item was lower than the chance level in the Tasty–Healthy trials [t(48) = −2.25; P < .05], but higher than the chance level in the Not tasty–Unhealthy trials [t(48) = 13.6; P < .001] (Fig. 2A, blue bars), suggesting that participants correctly understood the task and chose the food according to the individual preferences measured in the rating task (Fig. 1A).

**Figure 2.**
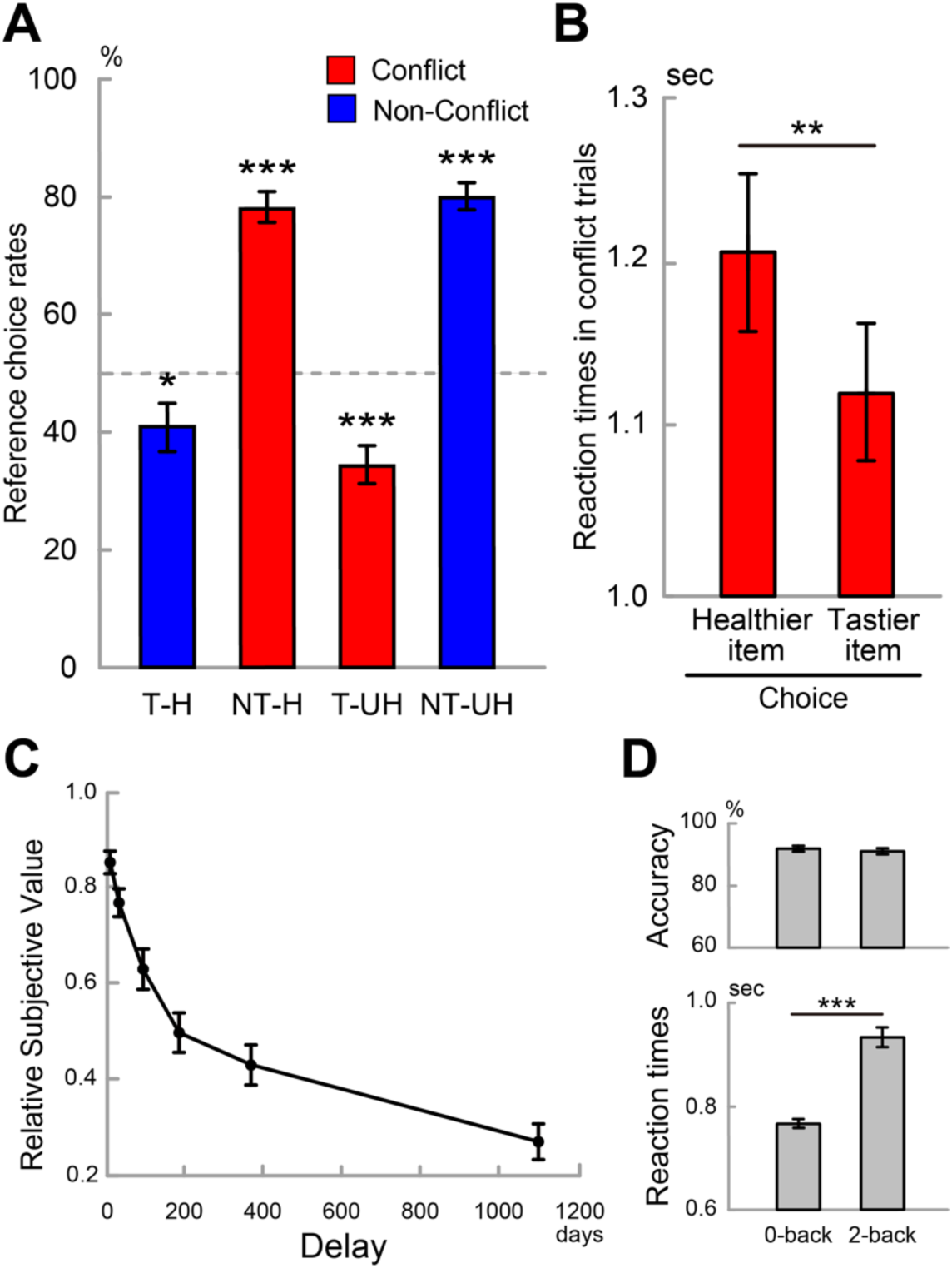
Behavioral results. ***A*,** Choice preference for the food choice task. The vertical axis indicates the percentage of trials in which the reference items were chosen, and the horizontal axis indicates trial types. The dotted line shows the chance level. T–H: Tasty–Healthy trial; NT–H: Not tasty–Healthy trial; T–UH: Tasty–Unhealthy trial; NT–UH: Not tasty–Unhealthy trial. The bars for the conflict and non-conflict trials were colored red and blue, respectively. ***: P < .001; **: P < .01; *: P < .05 relative to the chance level. ***B*,** Reaction times (RTs) in the conflict trials. The left bar shows RTs for the trials in which healthier and not tastier items were chosen, and the right bar shows RTs for the trials in which tastier and unhealthier items were chosen. ***C*,** Estimated subjective values of a delayed reward as a function of the delay duration. The vertical axis indicates the subjective value of the delayed reward standardized by its reward amount (10000 yen), and a value of 1.0 indicates that the delayed reward is not time-discounted. Error bars indicate standard errors of the mean across participants. ***D*,** Behavioral results of the WM task. Accuracy (*left*); RTs (*right*).

We next examined participants’ choice behavior in the conflict trials (Not tasty–Healthy and Tasty–Unhealthy trials), where either taste or healthiness was higher compared to reference items while the alternate property was lower. In the conflict trials, the rate of choosing the reference item was higher than the chance level in the Not tasty–Healthy trials and lower than the chance level in the Tasty–Unhealthy trials [Not tasty–Healthy: t(48) = 11.6; P < .001; Tasty–Unhealthy: t(48) = −4.9; P < .001](Fig. 2A, red bars). The choice preference in the conflict trials indicated that participants preferred tastier–unhealthier items rather than non-tastier–healthier items, suggesting that they prioritized taste over healthiness, consistent with prior studies (Hare et al., 2009, 2011).

Interestingly, in the conflict trials, reaction times (RTs) were longer when choosing healthier items (i.e., reference items in the Tasty–Unhealthy trials and target items in the Not tasty–Healthy trials) than when choosing tastier items (i.e., reference items in the Not tasty–Healthy trials and target items in the Tasty–Unhealthy trials) [t(48) = 2.7; P < .01] (Fig. 2B). Together, the behavioral results suggest that participants prioritized taste overall, but when choosing non-tasty–healthy foods over tasty– unhealthy foods, decision-making became difficult.

### Intertemporal decision-making task

Figure 2C shows the subjective value of the delayed monetary reward as a function of delay duration. Participants showed significant delay discounting [t(48) = 15.6; P < .001] and its degree became greater as the delay became longer [t(48) = −13.9; P < .001 with linear contrast] (Fig. 2C), reproducing the results of previous studies on delay discounting (Rachlin et al. 1991; Estle et al. 2007; Jimura *et al*. 2011; Jimura *et al*. 2018).

### WM task

Accuracy did not differ between the 2-back and 0-back conditions [t(48) = −0.8, P = 0.39], and RTs were longer in the 2-back condition than in the 0-back condition [t(48) = 12.4, P < .001] (Fig. 2D), suggesting that participants performed the task well but the 2-back condition required greater cognitive control demand, consistent with prior studies (Miller et al. 2009; Shucard et al. 2011).

### Imaging results

#### Food choice task

We first explored brain regions involved in prioritizing healthiness over taste, using data from the conflict trials of the food choice task. Figure 3 shows brain regions that exhibited greater activity when participants chose healthy items over tasty items in the Tasty–Unhealthy and Not tasty–Healthy trials. Greater activity was observed in multiple prefrontal regions, including the lateral and medial superior frontal cortex (SFC), dlPFC, anterior cingulate cortex (ACC), anterior insula (aINS), and vmPFC. A full list of coordinates is shown in Table S1. Notably, behavioral analysis showed prolonged RTs when participants prioritized healthiness (Fig. 2B), and these increased RTs were associated with the greater activity in these prefrontal regions. On the other hand, when choosing tasty items in the conflict trials, greater activity was observed in posterior brain regions, including the occipitotemporal cortex and cerebellar cortex (Lobules VI, VIIb, VIIIa, and VIIIb, and Crus I).

**Figure 3.**
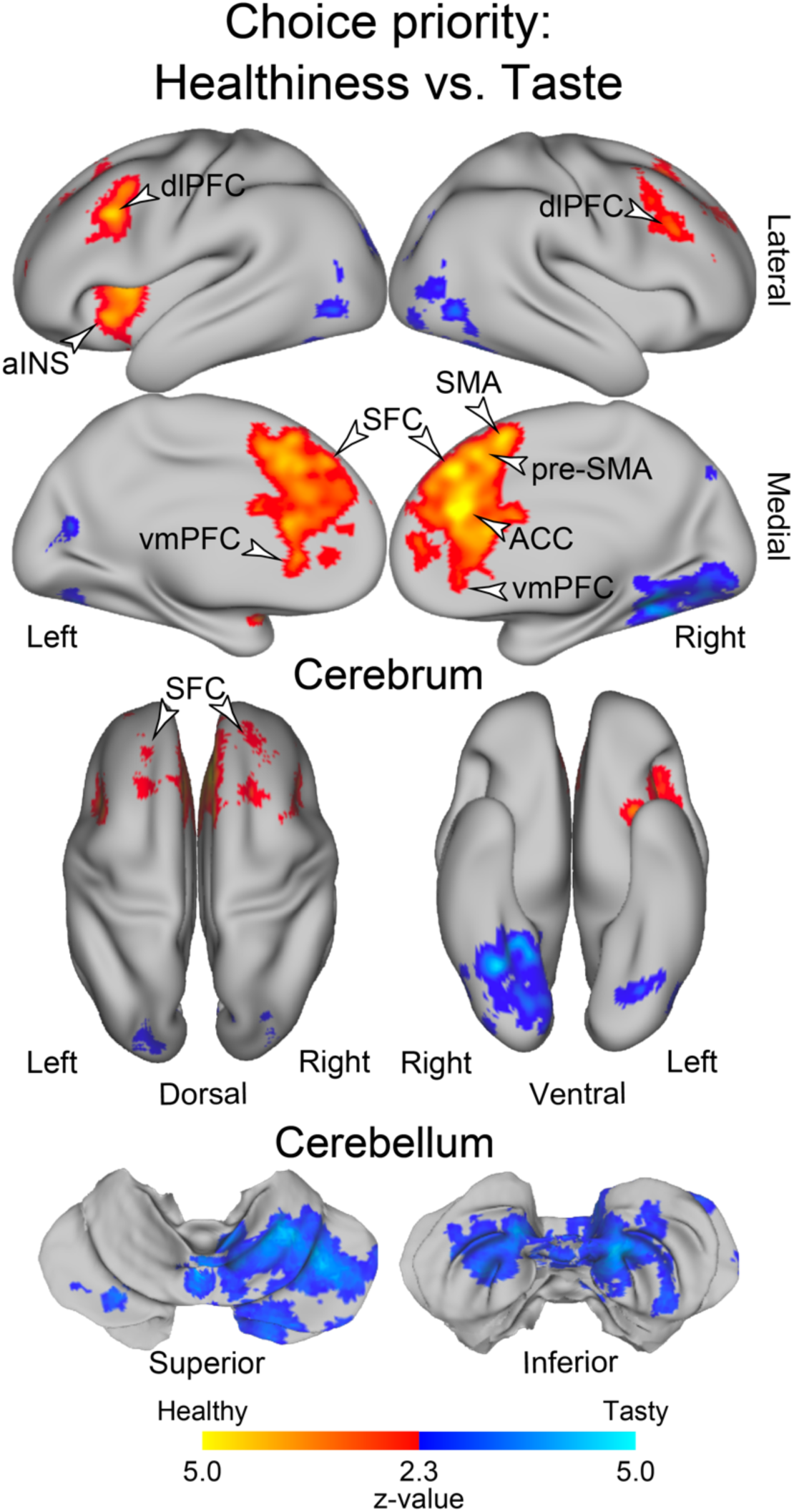
Statistical activation maps of the contrast for the choice behavior in the conflict trials where healthiness was prioritized over taste and where taste was prioritized over healthiness in the conflict trials (P < .05, FWE-corrected across the whole brain based on non-parametric permutation tests). Hot and cool colors indicate greater priority for healthiness and taste, respectively. Maps are overlaid onto a 3D surface of the brain. *Top*: Lateral medial surfaces of the cerebrum. *Middle*: Dorsal and ventral surfaces of the cerebrum. *Bottom*: Superior and inferior surfaces of the cerebellum. The white arrowheads indicate the superior frontal cortex (SFC), dorsolateral prefrontal cortex (dlPFC), anterior cingulate cortex (ACC), anterior insula (aINS), and ventromedial prefrontal cortex (vmPFC).

We next asked whether healthy food choice involves self-control. To this end, we examined whether the greater brain activity when choosing healthy items was modulated by the degree of delay discounting that is known to be linked to self-control in decision-making. Specifically, we explored brain regions that showed significant inter-participant correlations between the brain activity that occurred when choosing healthy items and AuC measures of the delay-discounting curve (see the “Statistical analysis” section in the Materials and Methods). A significant positive correlation was observed in multiple prefrontal regions, including the lateral and medial SFC, dlPFC, ACC, aINS and vmPFC (Fig. 4A and Table S2). Interestingly, these regions overlapped well with those showing significant mean positive activity when choosing healthy items (Fig. 3).

**Figure 4.**
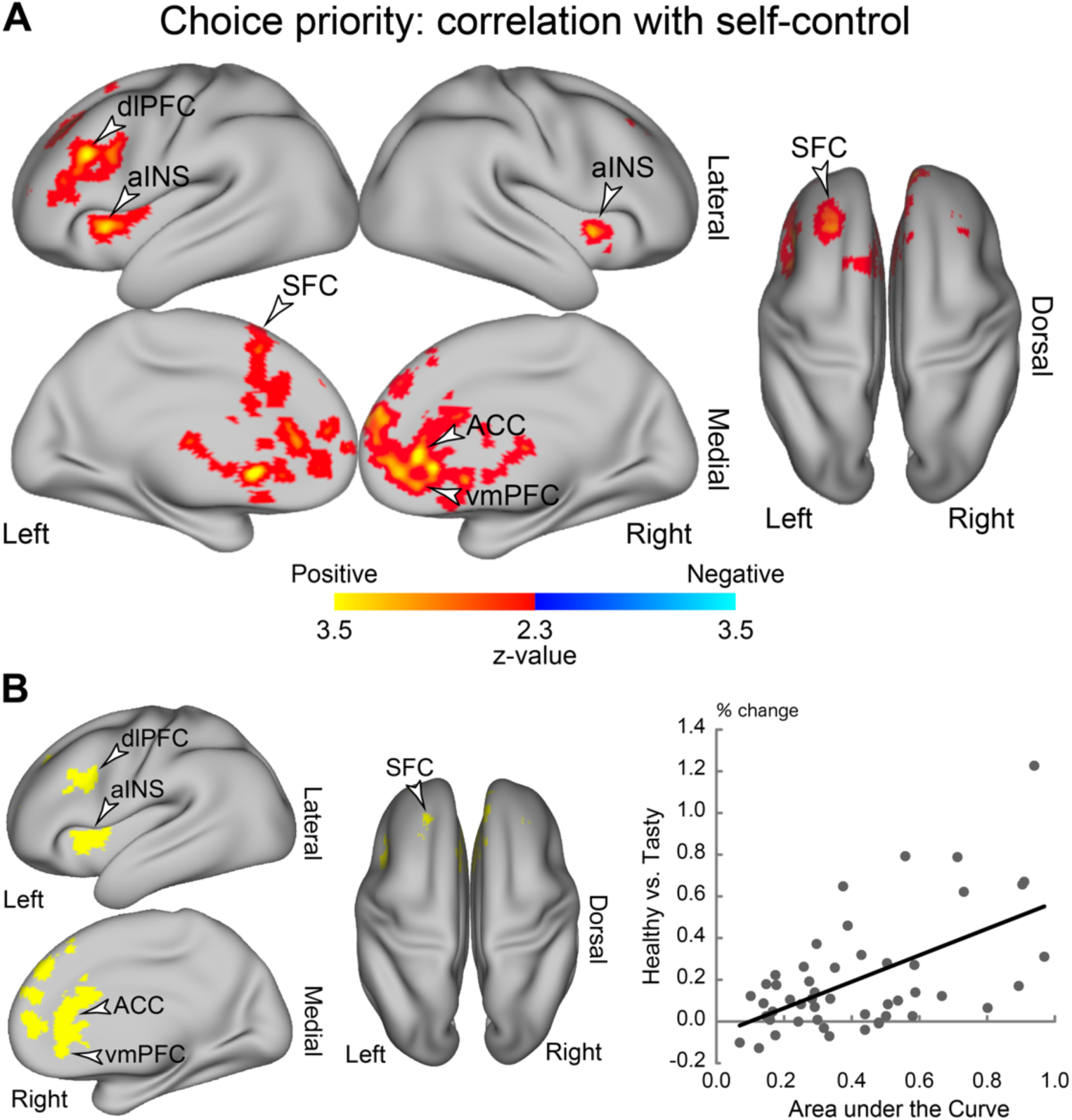
***A***, Statistical maps of the cross-subject correlation between the choice priority activity (Fig. 3) and the degree of self-control measured as subjective values of the delayed reward (AuC; Fig. 2C) (P < .05, FWE-corrected across the whole brain based on non-parametric permutation tests). Hot and cool colors indicate positive and negative correlations, respectively. Other formatting is similar to that in Fig. 3. ***B***, Conjunction analysis. The yellow areas (*left*) indicate the regions showing both significant positive activity while prioritizing healthiness (Fig. 3) and significant positive correlation (panel A). *Right*: A scatter plot of brain activity for the choice priority (healthiness vs. taste) against the degree of self-control (AuC) for the regions showing the conjunction effect in the left panel (yellow). The plots are for display purposes only, to avoid circular analysis. Each dot indicates one participant.

Then, to examine the spatial overlap of the maps of mean activity (Fig. 3) and the correlation between this activity and the AuC (Fig. 4A), we performed conjunction analysis. Conjunction effects were observed in multiple prefrontal regions (Fig. 4B *left* and Table S2). A scatter plot in Figure 4B shows an example the conjunction effect (only for display purposes to avoid circular analysis). The results of the conjunction analysis indicate that the lateral and medial SFC, dlPFC, ACC, aINS, and vmPFC are involved in choosing healthy foods, and the degree of their involvements is greater in high self-controlled individuals.

### WM task

Figure 5A shows activation maps for the contrast 2-back vs. 0-back conditions. Consistent with well-established results in previous studies, strong activation was observed in fronto-parietal regions, the media prefrontal cortex, ACC, superior cerebellar cortex (lobule VI and crus I), and inferior cerebellar cortex (lobule VIIb and Crus II) (Miller and Cohen 2001; Ragland et al. 2002; Botvinick 2007; Kim et al. 2012; Guell et al. 2018). These regions are also well known to implement core mechanisms of cognitive control (Okayasu et al. 2023).

**Figure 5.**
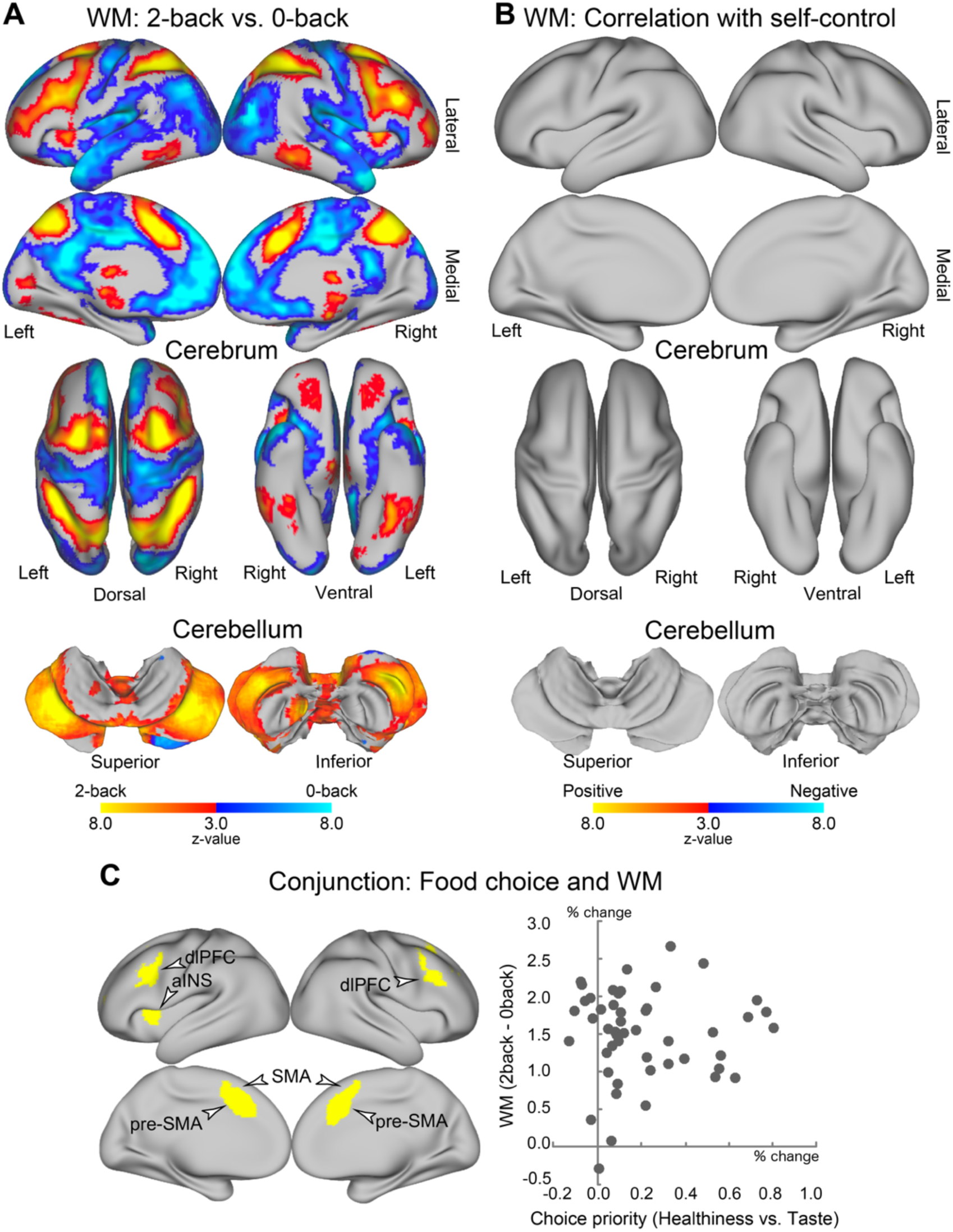
***A***, Statistical activation maps of the working memory (WM) load effect in N-back tasks (2-back vs. 0-back) (P < .05, FWE-corrected across the whole brain based on non-parametric permutation tests). Hot and cool colors indicate greater activity in the 2-back and 0-back tasks, respectively. Other formatting is similar to those in Fig. 3. ***B***, Statistical maps of the correlation between brain activity for the of WM load effect and the degree of self-control (AuC) (P < .05, FWE-corrected across the whole brain based on non-parametric permutation tests). Other formatting is similar to that in Fig. 4A. ***C***, Conjunction analysis of the food choice task and WM task (*left*). The yellow areas indicate the regions showing significant positive activity for both the choice priority effect (Fig. 3) and the WM load effect (Fig. 4A). A scatter plot of brain activity for the WM load effect against the choice priority effect (*right*). Each dot indicates one participant.

Previous behavioral and neuroimaging studies have suggested that WM mechanisms guide self-control in intertemporal choice (Hinson *et al*. 2003; Whitney et al. 2004; Shamosh *et al*. 2008; Jimura *et al*. 2018). To test the hypothesis that WM mechanisms are associated with the degree of delay discounting, we explored correlations between WM activity (2-back vs. 0-back) and AuC measures across participants. Surprisingly, significant correlations were absent across the whole brain (Fig. 5B), which were not compatible with the previous neuroimaging and behavioral studies (Hinson *et al*. 2003; Whitney *et al*. 2004; Shamosh *et al*. 2008; Jimura *et al*. 2018; Matsui *et al*. 2022)

If self-control in food choice is guided by cognitive control, common cognitive and decision processing should occur in both the food choice and WM tasks because the WM task requires cognitive control. In particular, 1) these two tasks should activate common brain regions and 2) the activation in the overlapping regions should be correlated between the two tasks (Jimura *et al*. 2018). To test the former hypothesis, we first performed a conjunction analysis for the food choice task (healthy items against tasty items; Fig. 3) and WM task (2-back vs. 0-back; Fig. 5A). The analysis revealed a joint effect in multiple lateral and medial prefrontal cortices, including the dlPFC, pre-SMA, SMA, and aINS (Fig 5C *left*). We then tested the latter hypothesis by examining a cross-subject correlation between WM activity and healthy food choice activity in the region exhibiting the joint effect. The analysis revealed no correlation [r = .022, t(47) = 0.15, P = .88; Fig. 5C *right*]. This lack of correlation suggests that decision and cognitive processing in these regions is independent despite the spatial overlap of brain activity. This phenomenon has been described as multi-independent processing (Poldrack 2006; Fox and Friston 2012) and the multiple-demand system (Duncan 2010; Camilleri et al. 2018) (see Discussion for details).

### Distinct prefrontal regions involved in healthy food choice

We have so far identified two types of brain regions showing prominent activity during healthy food choice: 1) regions whose activity becomes greater in individuals with smaller delay discounting (Fig. 4B) and 2) regions also involved in WM but whose activity is not correlated between WM and healthy food choice (Fig. 5C).

To further characterize these brain regions, we performed a conjunction analysis of the two types of the regions. Figure 6 shows conjunction and disjunction of the two maps (Fig 4B *left* and Fig. 5C *left*). The dlPFC, pre-SMA, SMA, and aINS show a single (i.e., disjunction) effect of WM and food choice only (blue regions in Fig. 6) or a conjunction effect of the two (cyan regions in Fig. 6). This indicated that these regions were activated during both WM and food choice tasks, but their activity during healthy food choice was unrelated to WM.

**Figure 6.**
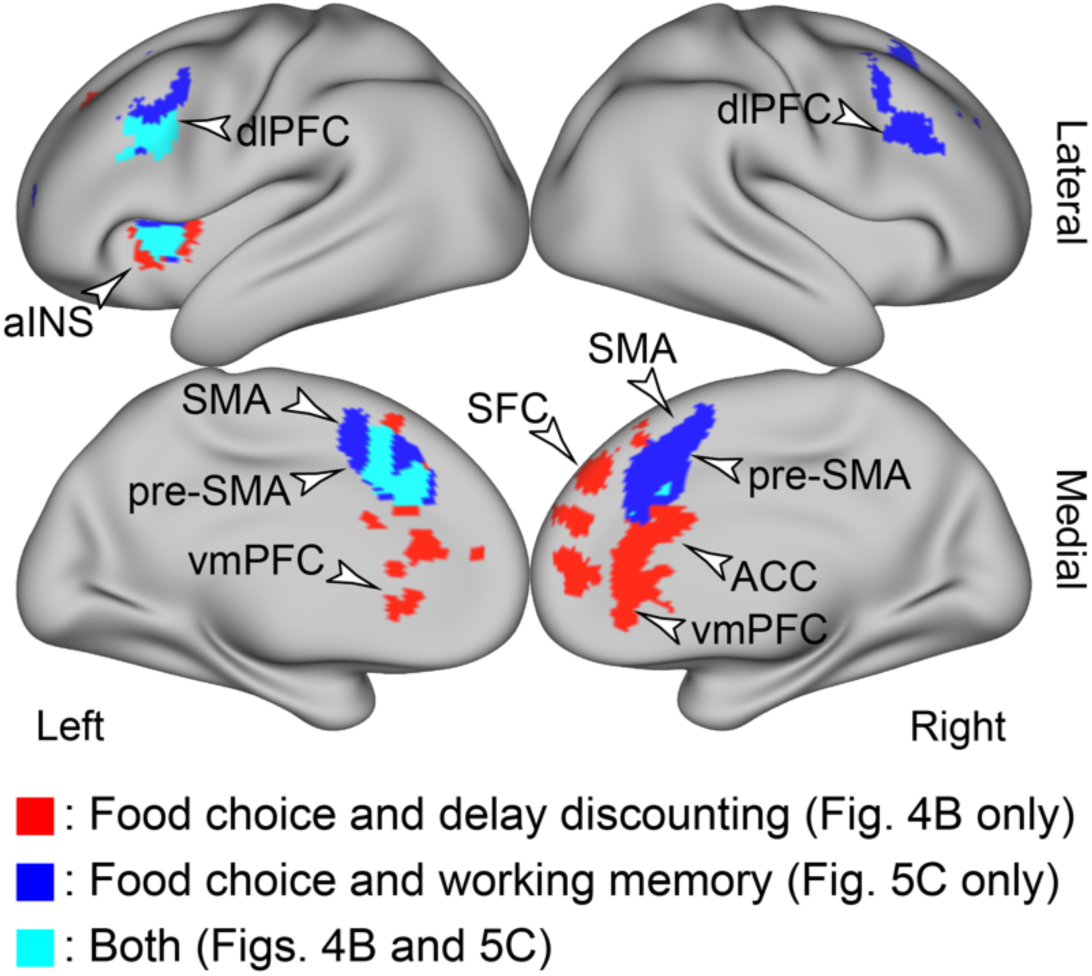
***A*,** The maps show conjunction and disjunction of brain regions exhibiting a conjoint effect of the food choice task and correlation with delay discounting (Fig. 4B), and also a conjoint effect of food choice task and the working memory task (Fig. 5C). Red: regions showing a conjoint effect of food choice and delay discounting only (Fig. 4B only); blue: regions showing a conjoint effect of food choice and working memory only (Fig. 5C only); cyan: regions showing both of the conjoint effects (Figs. 4B and 5C).

On the other hand, the activity in the SFC, vmPFC and ACC showed a single (i.e., disjunction) effect of the correlation between healthy food choice activity and delay discounting only (red regions in Fig. 6). This indicated that their activity during healthy food choice was elevated in individuals with smaller delay discounting, but the regions were not involved in WM. Collectively, the results suggest that the SFC, vmPFC, and ACC play a role specific to healthy food choice without engaging WM.

### HCP data analysis

One potential concern is that the absence of a correlation between WM and food choice task is due to the small sample size (N = 49). To circumvent this issue, we based the WM task and self-control measure in the current study on the procedures used in the HCP, enabling supplementary analysis with a large sample size (N = 1075).

In the HCP, the WM activity (2-back vs. 0-back) showed spatial patterns (Fig. S1A) similar to those in our data (Fig. 5A). Next we explored the correlation between WM activity and the AuC of delay discounting. A region in the inferior frontal junction (IFJ) showed a significant positive correlation (Fig. S1B), indicating greater WM activity in high self-control individuals. This IFJ region is located in a posterior part of the junction between the precentral sulcus (PCS) and the inferior frontal sulcus (IFS) (BA 6) (Fig. S1B *right*).

On the other hand, in the food choice task of the current study, the prefrontal regions showing correlations between food choice activity and the AuC were located in an anterior part of the IFJ (BA 6/44) and in the inferior frontal sulcus (BA 44/45) (Fig. S1C). The spatial segregation is clearly demonstrated by a map showing both types of correlations (Fig. S1D).

These collective results suggest that self-control is linked to cognitive control function, but this link is not associated with self-controlled food choice.

## Discussion

The current study examined the neural mechanisms underlying healthy food choices, and their relation to cognitive control–guided self-control. The lateral and medial SFC, dlPFC, vmPFC, ACC, and aINS were involved in prioritizing food healthiness over taste, and their activity was greater in individuals with high self-control who exhibited smaller delay discounting of a future monetary reward. On the other hand, WM activity was not associated with self-control or food choice activity. Notably, the SFC, vmPFC, and ACC were specifically involved in healthy food choice. The analysis of HCP data revealed that WM activity in the IFJ was positively correlated with self-control, but the IFJ region was spatially segregated from prefrontal regions showing correlations between self-control and activity during healthy food choice. These collective results suggest that healthy food choice involves prefrontal mechanisms that make it possible to maximize long-term rewards, not achieve cognitive control.

Choosing foods often imposes a tradeoff between taste and healthiness (Hare et al., 2011; Blechert et al., 2016., van der Laan et al., 2014). Notably, taste provides immediate satisfaction whereas healthiness promotes long-term welfare (Fishbach *et al*. 2003). This tradeoff involves a decision-making structure similar to the one involved in intertemporal decision-making (Hare et al. 2014; Zhang *et al*. 2022). Thus, prioritizing healthiness over taste may include a preference for a larger, delayed reward over a smaller, sooner reward. Self-control in intertemporal choice (i.e., the preference for a larger, delayed reward) and in food choice (i.e., prioritizing healthiness) may be explained by one unitary construct. Our results demonstrating correlations between prefrontal activity while prioritizing healthiness and the AuC of delay discounting support this hypothesis (Hare et al., 2009, 2011; Maier et al., 201).

The intake of high-calorie, tasty foods is strongly associated with obesity (Berthoud and Zheng 2012; Spetter *et al*. 2017), which suggests that obese subjects often choose impulsive behaviors that prioritize taste over healthiness (Davis *et al*. 2004; Schiff *et al*. 2016; Zhang *et al*. 2022). Indeed, previous studies demonstrated that obese subjects showed greater monetary delay discounting, indicating a preference for smaller immediate rewards over larger future rewards (Davis *et al*. 2004; Davis *et al*. 2010; Stoeckel *et al*. 2013; Lawyer *et al*. 2015; Zhang *et al*. 2022). Thus, the prefrontal regions that the current study identified as being involved in maximizing long-term reward may play a crucial role in resolving obesity.

Our imaging results demonstrated that dlPFC and vmPFC involvement in healthy food choice was associated with the degree of self-control in intertemporal choice of future monetary rewards. The dlPFC and vmPFC are known as crucial regions for self-controlled food choices (Hare et al., 2009, 2011; Maier et al., 2015; Spetter et al., 2017). It has been shown that in food choice, the dlPFC modulates the value signal represented in the vmPFC, which guides self-controlled food choice (Hare et al., 2009). Interestingly, intertemporal decision-making is also known to involve the dlPFC and vmPFC (McClure et al., 2004, 2007; Kable and Glimcher, 2007; Figner et al., 2010).

Specfically, the dlPFC guides self-controlled choice (McClure et al., 2004, 2007; Figner et al., 2010; Jimura et al., 2018) whereas the vmPFC respresents future reward values (Kable and Glimcher, 2007; Jimura et al., 2013). Delay discounting correlates with functional connectivity between the dlPFC and vmPFC during intertemporal choice (Hare et al., 2014). These results regarding food choice and intertemporal choice collectively suggest common prefrontal mechanisms of self-control. Our results are the first to demonstrate empirical evidence for their relationships. Importantly, our intertemporal decision-making task used a standard delay duration (3 years at maximum), which may be comparable to a time scale for consideration of health benefits. Taken together, dlPFC and vmPFC involvement in healthy food choices may recruit neural mechanisms that maximize long-term rewards in intertemporal decision-making.

The dlPFC, pre-SMA, SMA, and aINS showed a conjunction effect of healthy food choice and WM, but their activity was uncorrelated between the two tasks (Fig. 5C). The absence of a correlation suggests that these prefrontal regions are involved in both tasks, but function independently in processing healthy food choice and WM. This phenomenon is known as multi-independent processing, where one brain region implements multiple distinct functions (Poldrack 2006; Fox and Friston 2012). It is often observed in association cortices but not so often in sensory and motor cortices, and is one important source of the difficulty in reverse inference in association cortices (Yarkoni et al. 2011). These prefrontal regions mentioned above are also involved in the multiple-demand system that implements a wide range of cognitive functions (Duncan 2010; Camilleri *et al*. 2018; Tsumura et al. 2021; Tsumura et al. 2022).

In contrast, the SFC, ACC, vmPFC were not involved in WM, but their activity during healthy food choice was increased in individuals with smaller delay discounting (Fig. 6). The vmPFC and rostral portion of the ACC are known to implement the value-related systems (Kable and Glimcher 2007; Levy and Glimcher 2012) and are involved in healthy food choice (Hare et al., 2009, 2011; Maier et al., 2015; Spetter et al., 2017). Thus, an interesting finding of our study is the involvement of the lateral and medial parts of the SFC (BA 9 and 8) in healthy food choice (Fig. 3 and Table S1), which was enhanced in self-controlled individuals (Fig. 4 and Table S2). Previous neuroanatomical studies showed that the SFC cytoarchitectonically differs from the middle frontal gyrus that includes the dlPFC labeled as BA 9/46 (Petrides et al. 2012). It is thus possible that the role of the SFC in healthy food choice is distinct from those of the dlPFC and vmPFC. Further research is needed to clarify the function of the SFC, but our study suggests that it may cooperate with the dlPFC and vmPFC in choosing healthy foods.

Despite the strong relationship between self-control in food choice and intertemporal choice, our study showed that cognitive control and self-control were unrelated. Specifically, WM activity in our data was not correlated with the degree of self-control (Fig. 5B). To further explore the relationships between cognitive control and self-control, we analyzed large-sample data obtained from the HCP, and identified a statistically significant correlation in a posterior part of the IFJ (BA6), a well-known area involving cognitive control. Nonetheless, this correlation may reflect different functioning from that involved in food choice, as the correlation between food choice and self-control was identified more anteriorly (BA 6/44) to the IFJ correlation (Fig. 6D). Together, our study suggests that cognitive control may be weakly associated with self-control, possibly reflecting another aspect of self-control that is not linked to dietary choice. Confirmation of such a weak effect would require a large sample size.

A previous study showed that that WM activity in the prefrontal cortex reflects the degree of delay discounting and mediates the relationships between self-control and cognitive ability (Shamosh *et al*. 2008). Interestingly, cognitive ability is strongly associated with WM functions in the dlPFC (Gray *et al*. 2003), and the mediating role of the prefrontal cortex may be indirectly related to self-control.

Our previous study demonstrated that lateral prefrontal cortex activity was correlated between during a WM task and difficult decision-making in an intertemporal choice task, suggesting that difficult decision-making involves elaboration of choice-relevant information achieved by cognitive control (Jimura *et al*. 2018). A key component of the previous study is that we examined choice difficulty effects independently of intertemporal choice effects *per se* (i.e., self-control or impulsivity) because the choice effect was subtracted out. Another study showed that complex decision-making where a choice option involved multiple factors that formed high dimensionality of choice information involved fronto-parietal mechanisms associated with cognitive control (Matsui *et al*. 2022). As such, the involvement of cognitive control in value-based decision-making may reflect increased cognitive demands for computation of the choice information, but not self-control. The results of the current study also support this possibility. Thus, the involvement of the lateral prefrontal cortex may not be unique to intertemporal decision-making or food choice.

We examined the correlation of task-related brain activity between healthy food choices and cognitive control, to test the hypothesis that cognitive control guides self-control in food choice. The lateral and medial prefrontal cortices showed a joint effect, but the conjoint regions were activated independently during WM and healthy food choice (Fig. 5C), suggesting that these regions operated independently in the two tasks. Such brain regions implementing multiple, different functions have previously been identified mainly in association cortices (Poldrack 2006; Fox and Friston 2012). Our results suggest that healthy food choice can be achieved independently of WM.

## Acknowledgements

This study was supported by JSPS Kakenhi, 17K01989, and 20H00521, and 21H05060 to KJ and 17H06033 and 21H05060 to JC, and NIPS Cooperative Study Program (23-623, 22-628, 21-554, 20-639, 19-635, 18-633, and 17-6229) to KJ,.

## Supplementary Material

**Figure S1.**
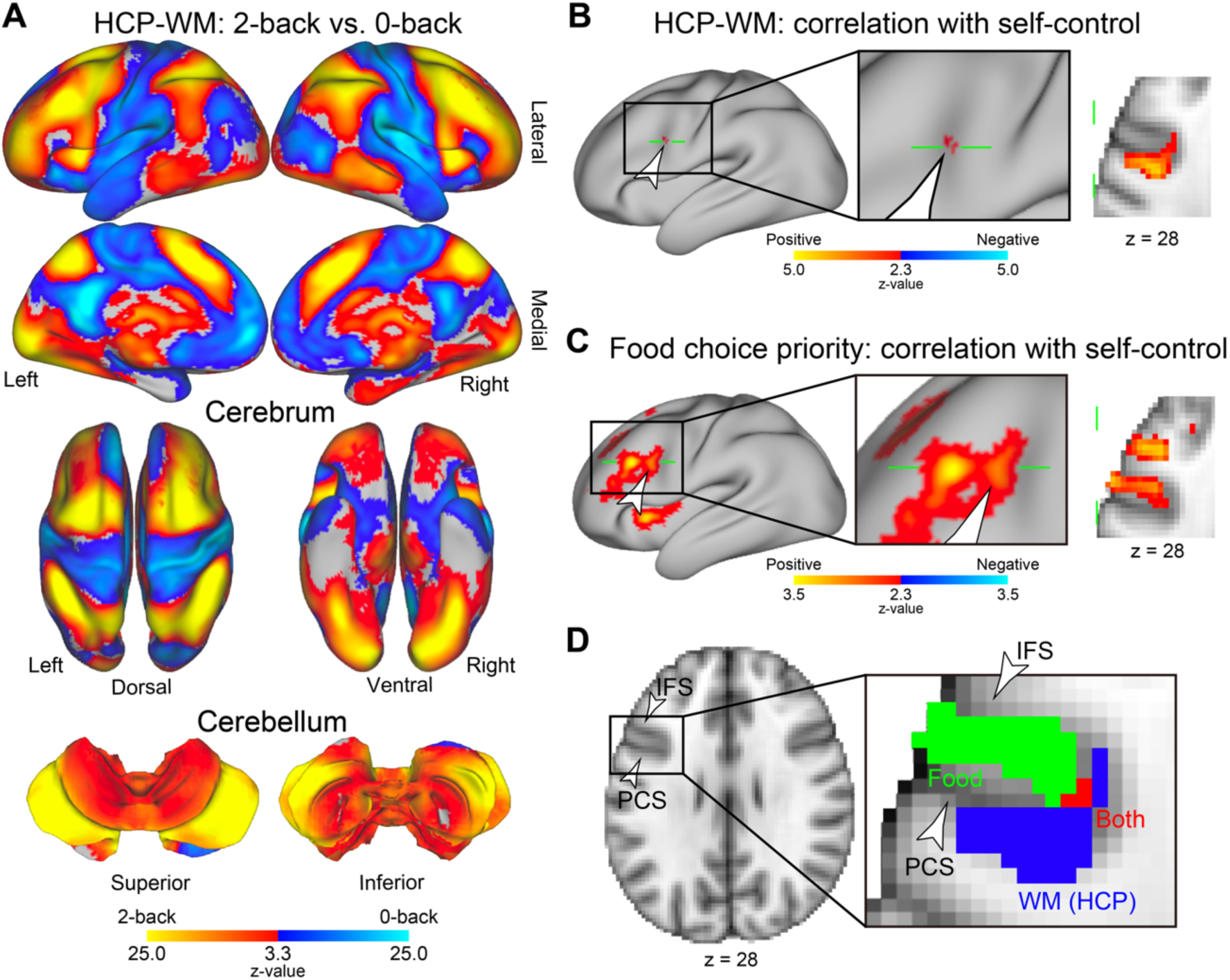
***A***, Statistical activation maps of the working memory (WM) load effect in the N-back task (2-back vs. 0-back) in the HCP data. Formatting is similar to that in Fig. 5A. ***B***, Statistical maps of the correlation between brain activity for the WM load effect and the degree of self-control (AuC) in the HCP data. The white arrowhead indicates a significant correlation in the inferior frontal junction. The area enclosed in the black rectangle (left) represents the area magnified in the middle. The green solid line on the surface indicates the axial section on the right. The level of the section is indicated at the bottom. ***C***, A map of correlations between the choice priority effect and self-control (Fig. 4A) is magnified at the inferior frontal area as in panel B. Formatting is similar to that in panel B. ***D***, The correlation effects in the food choice task (panel C) and the WM task (panel B) are mapped on a 2D section. The arrowheads indicate the inferior frontal sulcus (IFS) and precentral sulcus (PCS). The green and blue areas indicate the regions showing the correlation effect only in the food choice task and the WM task, respectively; the red area indicate the regions showing the correlation effect for both tasks. Other formatting is similar to that in panels B and C.

## Supplementary Tables

**Table S1.**
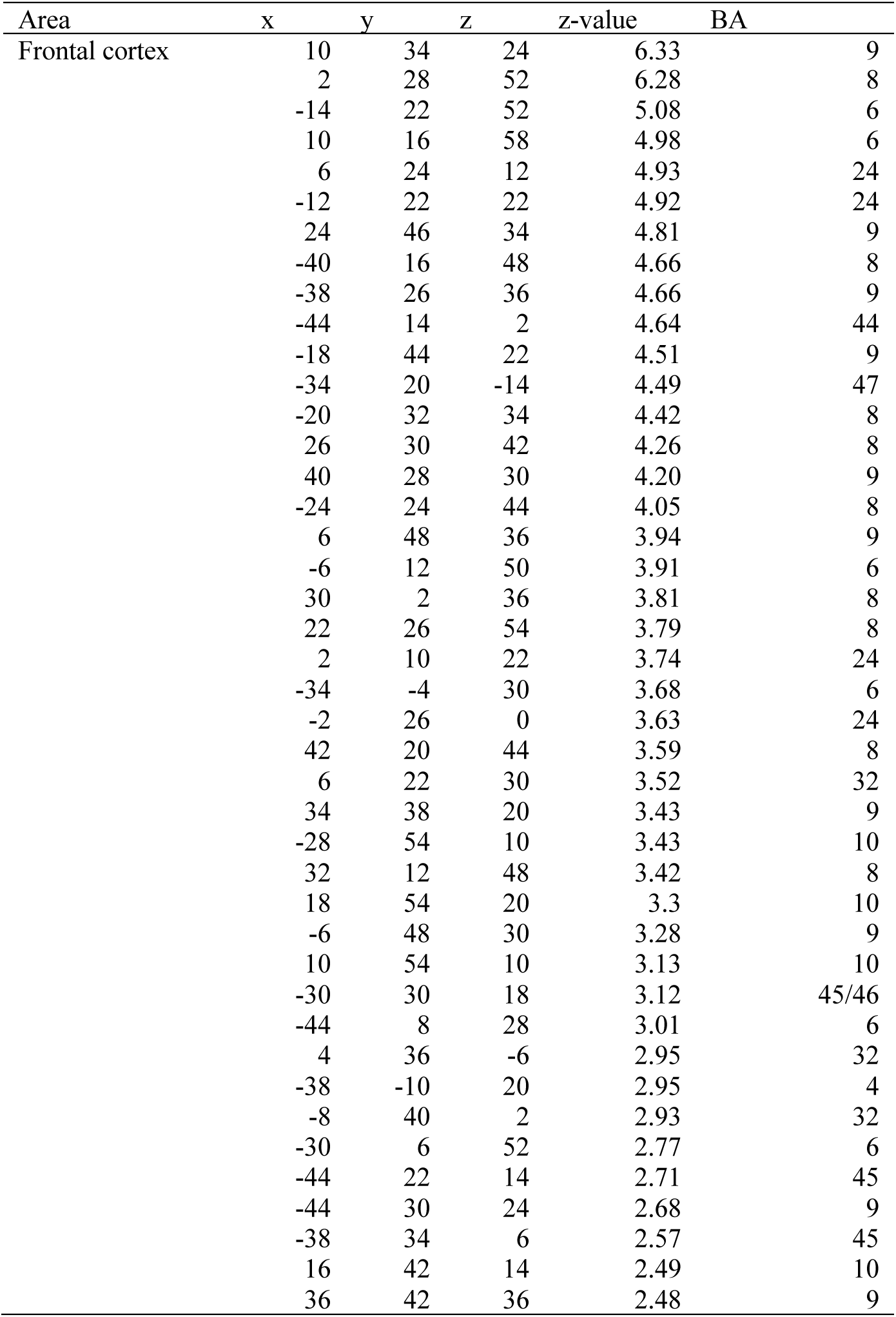
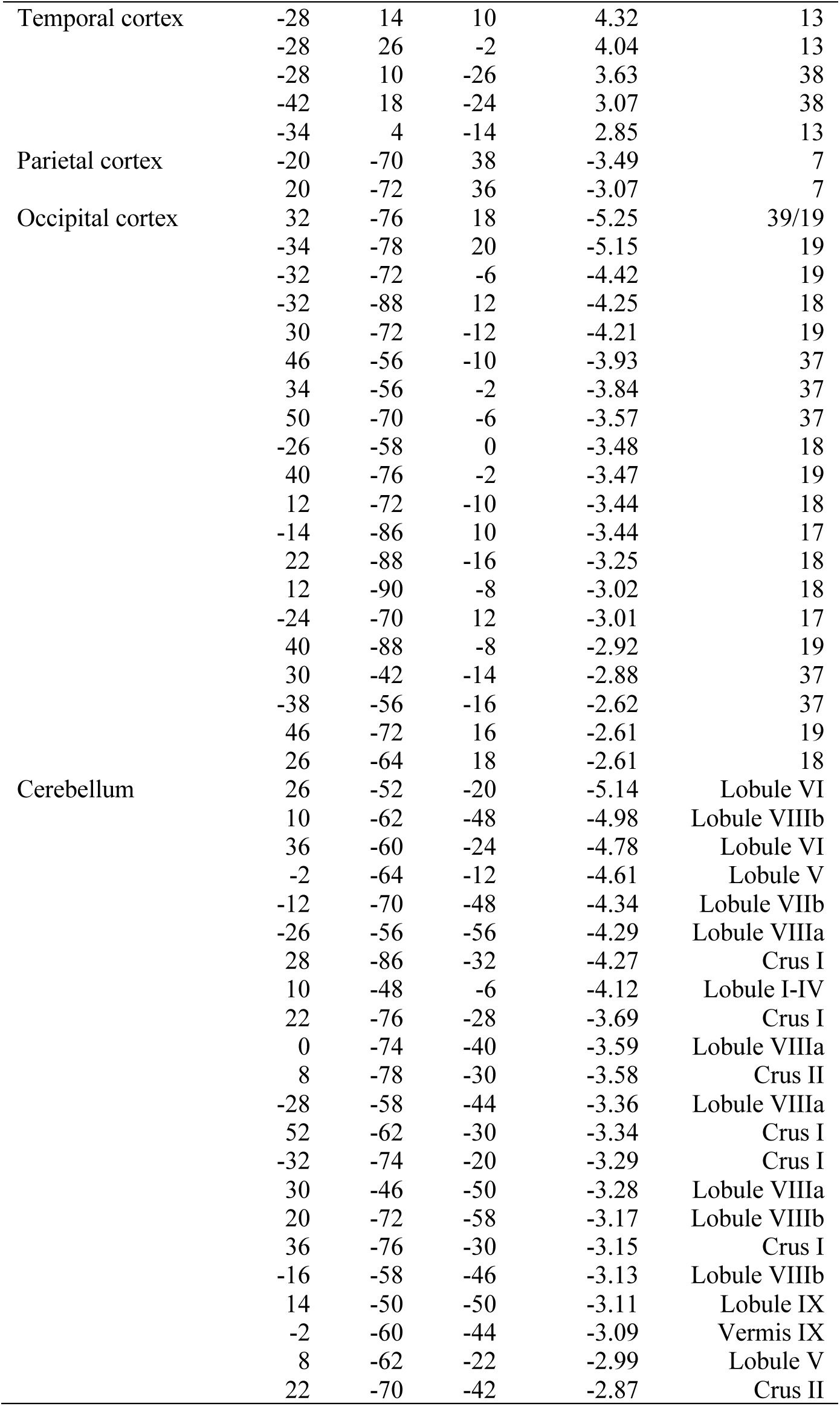
Brain regions showing a significant signal increase or decrease when choosing healthier and not tastier items in the conflict trials in the food choice task. Positive and negative z-values indicate greater signals in the trials with healthier or tastier items, respectively. Coordinates are listed in MNI space. BA indicates Brodmann area and is approximate.

**Table S2.**
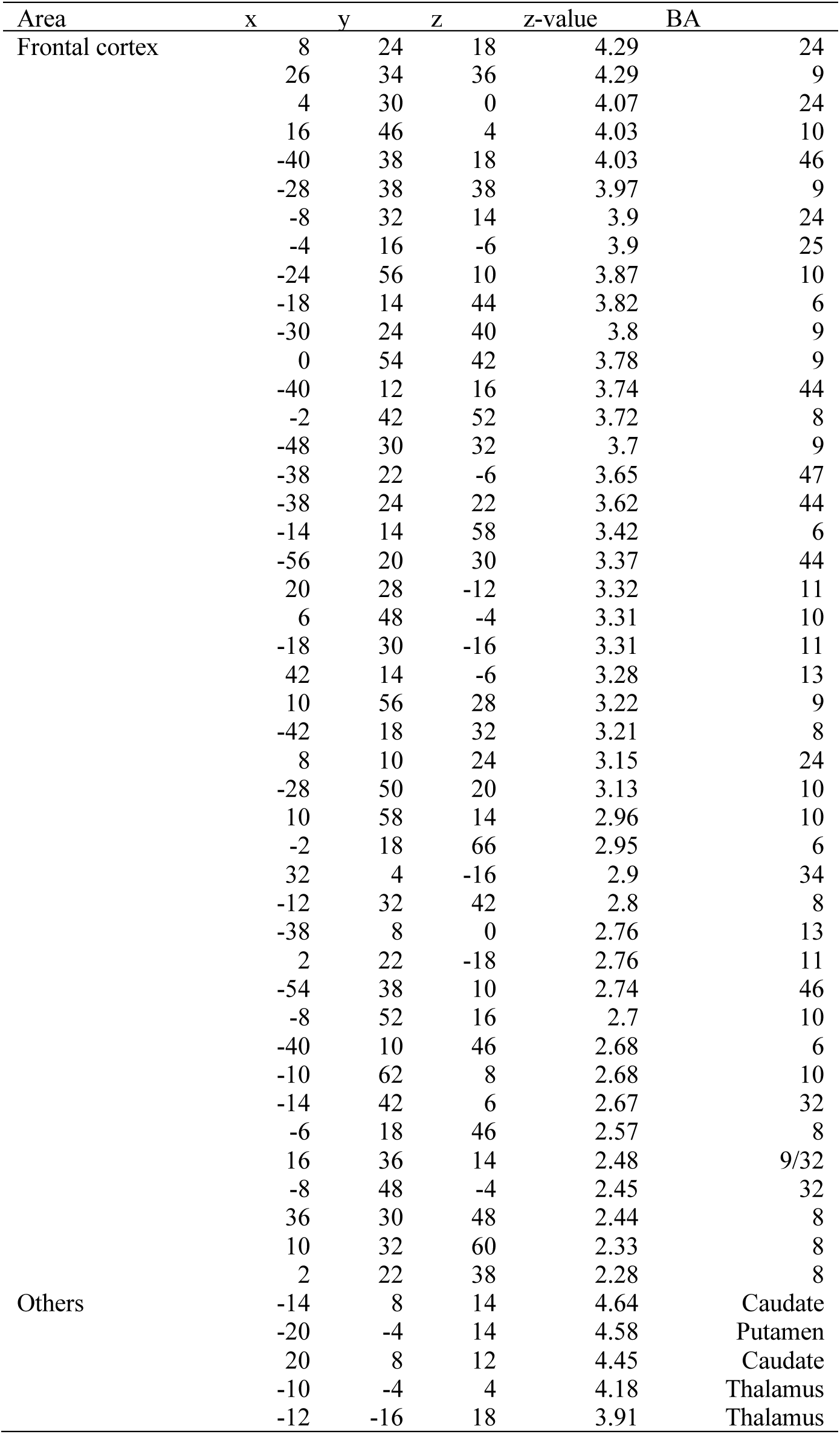
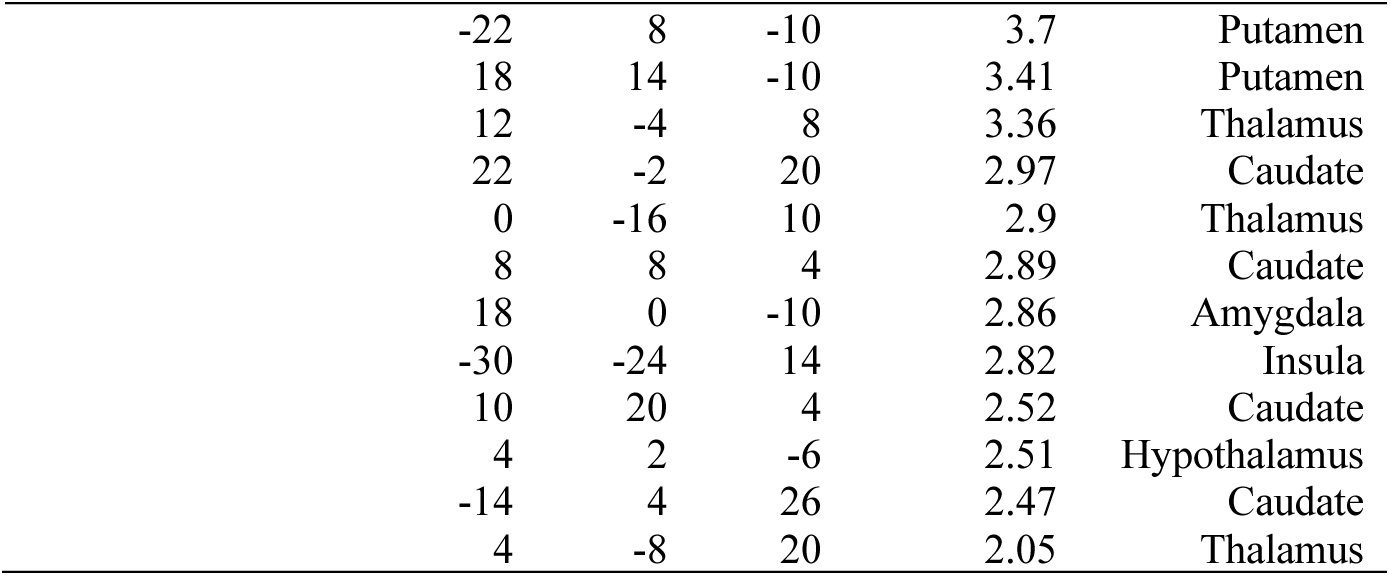
Brain regions showing significant correlation between brain activity when choosing healthier items and the degree of self-control. Formatting is similar to that in Table 1.

| Area | x | y | z | z-value | BA |
| --- | --- | --- | --- | --- | --- |
| Frontal cortex | 8 | 24 | 18 | 4.29 | 24 |
|  | 26 | 34 | 36 | 4.29 | 9 |
|  | 4 | 30 | 0 | 4.07 | 24 |
|  | 16 | 46 | 4 | 4.03 | 10 |
|  | -40 | 38 | 18 | 4.03 | 46 |
|  | -28 | 38 | 38 | 3.97 | 9 |
|  | -8 | 32 | 14 | 3.9 | 24 |
|  | -4 | 16 | -6 | 3.9 | 25 |
|  | -24 | 56 | 10 | 3.87 | 10 |
|  | -18 | 14 | 44 | 3.82 | 6 |
|  | -30 | 24 | 40 | 3.8 | 9 |
|  | 0 | 54 | 42 | 3.78 | 9 |
|  | -40 | 12 | 16 | 3.74 | 44 |
|  | -2 | 42 | 52 | 3.72 | 8 |
|  | -48 | 30 | 32 | 3.7 | 9 |
|  | -38 | 22 | -6 | 3.65 | 47 |
|  | -38 | 24 | 22 | 3.62 | 44 |
|  | -14 | 14 | 58 | 3.42 | 6 |
|  | -56 | 20 | 30 | 3.37 | 44 |
|  | 20 | 28 | -12 | 3.32 | 11 |
|  | 6 | 48 | -4 | 3.31 | 10 |
|  | -18 | 30 | -16 | 3.31 | 11 |
|  | 42 | 14 | -6 | 3.28 | 13 |
|  | 10 | 56 | 28 | 3.22 | 9 |
|  | -42 | 18 | 32 | 3.21 | 8 |
|  | 8 | 10 | 24 | 3.15 | 24 |
|  | -28 | 50 | 20 | 3.13 | 10 |
|  | 10 | 58 | 14 | 2.96 | 10 |
|  | -2 | 18 | 66 | 2.95 | 6 |
|  | 32 | 4 | -16 | 2.9 | 34 |
|  | -12 | 32 | 42 | 2.8 | 8 |
|  | -38 | 8 | 0 | 2.76 | 13 |
|  | 2 | 22 | -18 | 2.76 | 11 |
|  | -54 | 38 | 10 | 2.74 | 46 |
|  | -8 | 52 | 16 | 2.7 | 10 |
|  | -40 | 10 | 46 | 2.68 | 6 |
|  | -10 | 62 | 8 | 2.68 | 10 |
|  | -14 | 42 | 6 | 2.67 | 32 |
|  | -6 | 18 | 46 | 2.57 | 8 |
|  | 16 | 36 | 14 | 2.48 | 9/32 |
|  | -8 | 48 | -4 | 2.45 | 32 |
|  | 36 | 30 | 48 | 2.44 | 8 |
|  | 10 | 32 | 60 | 2.33 | 8 |
|  | 2 | 22 | 38 | 2.28 | 8 |
| Others | -14 | 8 | 14 | 4.64 | Caudate |
|  | -20 | -4 | 14 | 4.58 | Putamen |
|  | 20 | 8 | 12 | 4.45 | Caudate |
|  | -10 | -4 | 4 | 4.18 | Thalamus |
|  | -12 | -16 | 18 | 3.91 | Thalamus |

|  |  |  |  |  |
| --- | --- | --- | --- | --- |
| -22 | 8 | -10 | 3.7 | Putamen |
| 18 | 14 | -10 | 3.41 | Putamen |
| 12 | -4 | 8 | 3.36 | Thalamus |
| 22 | -2 | 20 | 2.97 | Caudate |
| 0 | -16 | 10 | 2.9 | Thalamus |
| 8 | 8 | 4 | 2.89 | Caudate |
| 18 | 0 | -10 | 2.86 | Amygdala |
| -30 | -24 | 14 | 2.82 | Insula |
| 10 | 20 | 4 | 2.52 | Caudate |
| 4 | 2 | -6 | 2.51 | Hypothalamus |
| -14 | 4 | 26 | 2.47 | Caudate |
| 4 | -8 | 20 | 2.05 | Thalamus |

